# Occipitotemporal Representations Reflect Individual Differences in Conceptual Knowledge

**DOI:** 10.1101/264895

**Authors:** K. Braunlich, B. C. Love

## Abstract

Through selective attention, decision-makers can learn to ignore behaviorally-irrelevant stimulus dimensions. This can improve learning and increase the perceptual discriminability of relevant stimulus information. To account for this effect, popular contemporary cognitive models of categorization typically include of attentional parameters, which provide information about the importance of each stimulus dimension in decision-making. The effect of these parameters on psychological representation is often described geometrically, such that perceptual differences over relevant psychological dimensions are accentuated (or stretched), and differences over irrelevant dimensions are down-weighted (or compressed). In sensory and association cortex, representations of stimulus features are known to covary with their behavioral relevance. Although this implies that neural representational space might closely resemble that hypothesized by formal categorization theory, to date, attentional effects in the brain have been demonstrated through powerful experimental manipulations (e.g., contrasts between relevant and irrelevant features). This approach sidesteps the role of idiosyncratic conceptual knowledge in guiding attention to useful information sources. To bridge this divide, we used formal categorization models, which were fit to behavioral data, to make inferences about the concepts and strategies used by individual participants during decision-making. We found that when greater attentional weight was devoted to a particular visual feature (e.g., “color”), its value (e.g., “red”) was more accurately decoded from occipitotemporal cortex. We additionally found that this effect was sufficiently sensitive to reflect individual differences in conceptual knowledge. The results indicate that occipitotemporal stimulus representations are embedded within a space closely resembling that proposed by classic categorization models.

## Introduction

Through selective attention, knowledge of abstract concepts can emphasize relevant stimulus features. For example, while the size of garments is critical when choosing what to purchase, weight may be more important when deciding how to ship them. The attention devoted to individual features is flexibly-modulated according to current goals, transient contextual demands, and evolving conceptual knowledge (Tversky, 1977; Goldstone, 2003). In formal theories of categorization, one way to account for this is through inclusion of attentional parameters, which reflect the influence of each dimension on the final choice (e.g., Kruschke, 1992; Nosofsky, 1986; Love et al., 2004). These parameters are often described as “warping” multidimensional psychological space, such that differences along relevant stimulus dimensions are accentuated (or “stretched”) and differences along irrelevant dimensions are down-weighted (or “compressed”; Figure 1). Here, we directly test this classic idea by investigating whether these attentional parameters predict the strength of neural stimulus feature representations. Importantly, we link individual differences in conceptual knowledge (as revealed by model fits of attentional parameters) to individual differences in neural representation (as revealed by decoding stimulus features in fMRI data). In doing so, we bridge behavioral and neural levels of analysis at the individual level using cognitive models.

**Figure 1:**
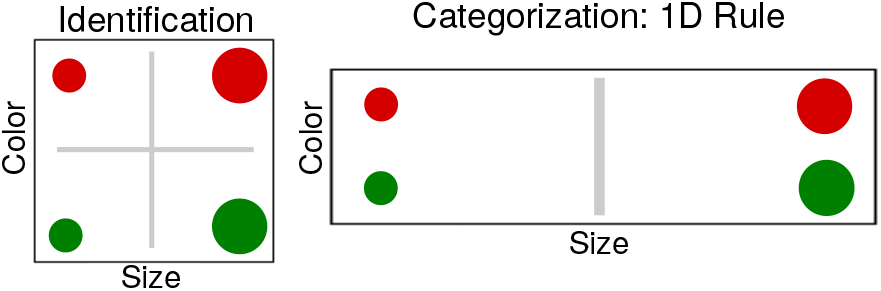
Example: Attention Influences Psychological Space. **Left**: In an object identification task, both psychological dimensions should receive equivalent attention, as they are equally relevant. **Right**: In a one-dimensional rule-based categorization task, only a single dimension is relevant (in this example, size), and decision-makers could ignore the irrelevant dimension (color). This is often described as “warping” psychological space such that differences along relevant dimensions are accentuated (or “stretched”), and differences along irrelevant dimensions are down-weighted (or “compressed”).

When identifying specific objects, agents must typically consider all stimulus features, and the psychological distance between stimuli closely reflects their perceptual attributes (Shepard, 1957; Townsend & Ashby, 1982). During categorization, however, groups of distinct stimuli must be treated equivalently, and both learning and generalization can be improved by selectively attending to relevant stimulus dimensions (Shepard et al., 1961; Nosofsky, 1986). Although categorization models differ in how stimuli are represented in memory (e.g., as individual exemplars, as prototypes, or as clusters that flexibly reflect environmental structure; Love et al., 2004; Nosofsky, 1987; Nosofsky & Zaki, 2002; Smith & Minda, 1998; Minda & Smith, 2002; Zaki et al., 2003), they assume that categorization involves learning to distribute attention across stimulus features so as to optimize behavioral performance. Although they differ in their mathematical details, these models also propose that *endogenous* (i.e., “top-down”) attentional control (Miller & Cohen, 2001; Tsotsos, 2011) can modulate the influence of the *exogenous* (or perceptual) stimulus dimensions on the behavioral choice. The attentional parameters play a key role in allowing the models to capture patterns of human generalization across different goals and rules. They also predict human eye-movements during category decision-making (e.g., Rehder & Hoffman, 2005a, b), and are therefore thought to reflect the strategies used by decision-makers to actively-sample information from the external world.

In the brain, effects of endogenous attention have been observed across the visual cortical hierarchy (Buffalo et al., 2010; Jehee et al., 2011; Kamitani & Tong, 2005, 2006; Luck et al., 1997; Motter, 1993). A general finding is that when attention is devoted to a specific visual feature, its neural representation is more accurately decoded. For instance, in human fMRI, when multiple visual gratings are concurrently presented, representations of attended orientations in areas V1-V4 are more easily decoded than those that are unattended (Kamitani & Tong, 2005; Jehee et al., 2011). Similarly, when random dot stimuli move in multiple directions, representations of attended motion directions in area MT+ are more easily decoded than those that are unattended (Kamitani & Tong, 2006). Whereas these studies have relied on explicit cues to guide attention to relevant aspects of the stimulus array, in real-world environments, decision-makers must typically rely on knowledge gained through past experience in order to selectively attend to relevant information sources.

Categorization tasks mirror this aspect of real-world environments; decision-makers must rely on learned conceptual knowledge in order to selectively attend to relevant stimulus dimensions. Several studies have investigated whether neural representations of exogenous information sources are modulated by learned conceptual knowledge (e.g., Li et al., 2007; Folstein et al., 2013; Sigala & Logothetis, 2002). Sigala & Logothetis (2002), for instance, trained macaques to categorize abstract images, which varied according to four stimulus dimensions. Neural representations of the two behaviorally-relevant stimulus dimensions (i.e., the dimensions that reliably predicted the correct response) in the inferior temporal lobe were enhanced relative to those of the irrelevant dimensions. Using fMRI with human participants, Li et al. (2007) investigated whether neural representations of stimulus motion and shape were influenced by their relevance to the active categorization rule. Using multivariate pattern analysis (MVPA), they similarly found that representations of these stimulus dimensions reflected their relevance to the active rule.

Across studies involving explicit attentional cues and categorization studies involving learned conceptual knowledge, a known effect is that occipitotemporal representations of behaviorally-relevant information sources are enhanced relative to those that are irrelevant (but this may not hold for integral stimulus dimensions; Garner, 1976). These effects are compelling, as they imply that occipitotemporal representational space may closely resemble that conceptualized by classic cognitive theory (e.g., Love et al., 2004; Kruschke, 1992; Nosofsky, 1986). Specifically, these representations may expand and contract, along axes defined by perceptually-separable stimulus dimensions (Garner, 1976), in ways that closely reflect the idiosyncratic concepts and strategies used by individual participants during decision-making.

Previous studies have relied on contrastive analyses in which neural representations of attended stimulus dimensions are compared to those of unattended dimensions. Although statistically-powerful, this approach defines selective attention in terms of the experimental paradigm (but see O’Bryan et al., 2018), and therefore sidesteps effects associated with individual differences in conceptual knowledge (e.g., Craig & Lewandowsky, 2012; Little & McDaniel, 2015; McDaniel et al., 2014; Raijmakers et al., 2014). These effects can be substantial, particularly for ill-defined categorization-problems (such as the 5/4 categorization task) that are common in every-day life (Hedge et al., 2017; Johansen & Palmeri, 2002). Here, we seek to address this divide by combining model-based fMRI (Turner et al., 2017; Palmeri et al., 2017) with multivariate pattern analyses. This allowed us to abstract away from individual differences in neural topography (Kriegeskorte & Kievit, 2013; Haxby et al., 2001; Haynes, 2015), to investigate whether neural stimulus representations reflect individual differences in conceptual knowledge. Specifically, we sought to investigate whether the attentional parameters derived from formal categorization models predict contortions of occipitotemporal representational space during decision-making.

We investigated this hypothesis using two publicly available datasets (osf.io). In the first (Mack et al., 2013), participants categorized abstract stimuli that varied according to four binary dimensions (Figure 2.A), according to a categorization strategy they learned prior to scanning. In the original paper, the authors fit both the Generalized Context Model (GCM; Nosofsky, 1986) and the Multiplicative Prototype Model (Nosofsky, 1987; Nosofsky & Zaki, 2002) to the behavioral data, and used them to compare exemplar and prototype accounts of occipitotemporal representation. Using representational similarity analysis (Kriegeskorte et al., 2008), Mack et al. (2013) additionally identified regions of the brain (lateral occipital cortex, parietal cortex, inferior frontal gyrus, and insular cortex) sensitive to the attentionally-modulated pairwise similarities between stimuli. Although these results (particularly those in lateral occipital cortex) imply that neural representations of the individual stimulus features might be modulated by selective attention, in principle, this could also reflect modulation within an abstract representational space where stimulus features are not individually represented. For instance, while visual cortex reflects sensory input (and is known to represent individual stimulus dimensions), prefrontal cortex can flexibly represent conjunctions of features, abstract rules, and category boundaries in a goal-directed manner. Representations in parietal cortex display intermediate characteristics, as they can reflect both sensory and decisional factors (Brincat et al., 2017; Jiang et al., 2007; Li et al., 2007).

**Figure 2:**
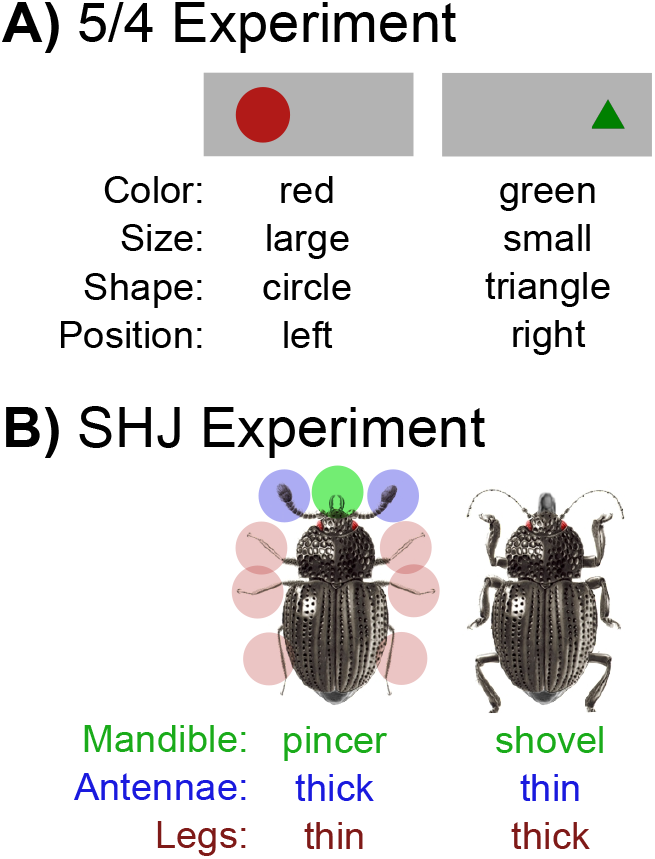
Stimuli. **A**) Two of the 16 stimuli used in the “5/4” experiment are illustrated. The stimuli varied according to four binary perceptual dimensions: color, size, shape and position. **B**) Two of the eight stimuli used in the SHJ experiment are illustrated. The stimuli were pictures of insects that varied according to three binary dimensions: mandible shape (highlighted in green), antennae thickness (highlighted in blue), and leg thickness (highlighted in red). For both experiments, the mapping of visual dimension to its role in each category structure (Tables 2 and 3) was randomized for each participant.

In the second dataset (Mack et al., 2016), participants learned, while scanning, to categorize images of insects that varied according to three binary perceptual dimensions (Figure 2.B), according to type I, type II and type VI problems described by Shepard et al. (1961).^1^ Importantly, although the same stimuli were included in each task, the degree to which each of the features predicted the correct choice differed between rules. The authors fit the SUSTAIN learning model (Supervised and Unsupervised STratified Adaptive Incremental Network; Love et al., 2004) to the behavioral data, and used it to investigate hippocampal involvement in the development of new conceptual knowledge. Using representational similarity analysis, they found that SUSTAIN successfully predicted the pairwise similarities between hippocampal stimulus representations across rule-switches. This suggests that hippocampal representations are updated according to goal-directed attentional selection of stimulus features.

## Methods

### Description of Datasets

In both experiments, participants categorized stimuli that were characterized by multiple perceptually-separable stimulus dimensions. As the mapping of perceptual attributes to their role in each category structure was randomized for each participant, it is possible to differentiate effects associated with intrinsic perceptual stimulus attributes from effects of behavioral relevance. For example, while color strongly predicted the correct category choice for some participants, it provided unreliable informative for others. In both experiments, participants were not instructed as to which cues were informative, and learned to perform each task through trial-and-error.

We used the GCM for the first dataset (the winning model from Mack et al. 2013), as participants learned how to perform the categorization task prior to scanning. We used SUSTAIN for the second dataset, as it learns on a trial-by-trial basis, and participants learned to perform each task during scanning. SUSTAIN was additionally fit in such a way that the learning of one task carried over to the next. Importantly, although the GCM and SUSTAIN differ in how stimuli are represented in memory (i.e., as exemplars or clusters), they similarly posit that attention “contorts” psychological space, as illustrated in Figure 1. Thus, these studies and models provide a good test of whether attention weights in successful cognitive models are plausible at both behavioral and neural levels of analysis.

### The “5/4” Dataset

The first dataset (Mack et al., 2013) was collected while 20 participants (14 Female) categorized abstract stimuli (Figure 2.A), which varied according to four binary stimulus dimensions (size: large vs. small, shape: circle vs. triangle, color: red vs. green, and position: left vs. right). Prior to scanning, they learned to categorize the stimuli according to the “5/4” categorization task (Medin & Schaffer, 1978) through trial-and-error. During this training session, participants were shown only the first nine stimuli shown in Table 2 (i.e., five category “A” members: A1-A5 and four category “B” members: B1-B4), and experienced 20 repetitions of each stimulus. During the anatomical scan, they additionally performed a “refresher” task, involving four additional repetitions on each training item. Each training trial involved a 3.5 second stimulus presentation period in which participants made a button press. Following the button press, a fixation cross was shown for 0.5 seconds, and feedback was then presented for 3.5 seconds. Feedback included information about the correct category, and about whether the response was correct or incorrect. During scanning, participants were required to categorize not only the training items, but also the seven transfer stimuli (i.e., T1-T7). In the scanner, stimuli were presented for 3.5 seconds on each trial, no feedback was provided, and stimuli were separated by a 6.5 second intertrial interval. Over six runs, each of the 16 stimuli were presented three times. The order of the stimulus presentations were randomized for each participant.

### The SHJ Dataset

In the second dataset (Mack et al., 2016), 23 right-handed participants (11 Female, mean age = 22.3 years) categorized images of insects (Figure 2.B) varying along three binary dimensions (legs: thick vs. thin, antennae: thick vs. thin, and mandible: pincer vs. shovel). We excluded data from two participants who each had corrupted data on one run. This resulted in 21 participants for the final analyses. During scanning, participants learned to categorize the stimuli according to the type I, type II, and type VI problems described by Shepard et al. (1961). In the type I problem, the optimal strategy required attending to a single stimulus dimension (e.g., “legs”) that perfectly predicted the category label, while ignoring the other two dimensions. In the type II problem, the optimal strategy was a logical XOR rule, in which two stimulus features had to be considered together. In the type VI problem, all stimulus features were relevant to the decision, and participants had to learn the mapping between individual stimuli and the category label. To maximally differentiate endogenous and exogenous factors, the irrelevant feature in the type II rule was used as a relevant feature of the type I problem for each participant.

Each problem was performed across four scanner runs. While all of the participants learned to perform the type VI problem first, the order of the type I and type II problems was then counterbalanced across participants. Each trial consisted of a 3.5 second stimulus presentation period, a jittered 0.5-4.5 second fixation period, and feedback. Feedback was presented for 2 seconds and consisted of an image of the presented insect, as well as text indicating whether the response was correct or incorrect. Each trial was separated by jittered intertrial interval (4-8 seconds), which consisting of a fixation cross. Each run included four presentations of each of the eight stimuli.

For consistency across datasets, we used the group-derived region of interest (ROI) used in “5/4” dataset (Figure 3.B), and performed a similar analysis. As participants in the SHJ experiment learned to perform the type I, type II and type VI problems during scanning, we mirrored the strategy used by the original authors, and divided the scanning sessions into early (first two runs of each problem) and late learning epochs (last two runs of each problem). We investigated the relationship between occipitotemporal representation and attention only during this late learning phase, in which behavior had largely stabilized.

**Figure 3:**
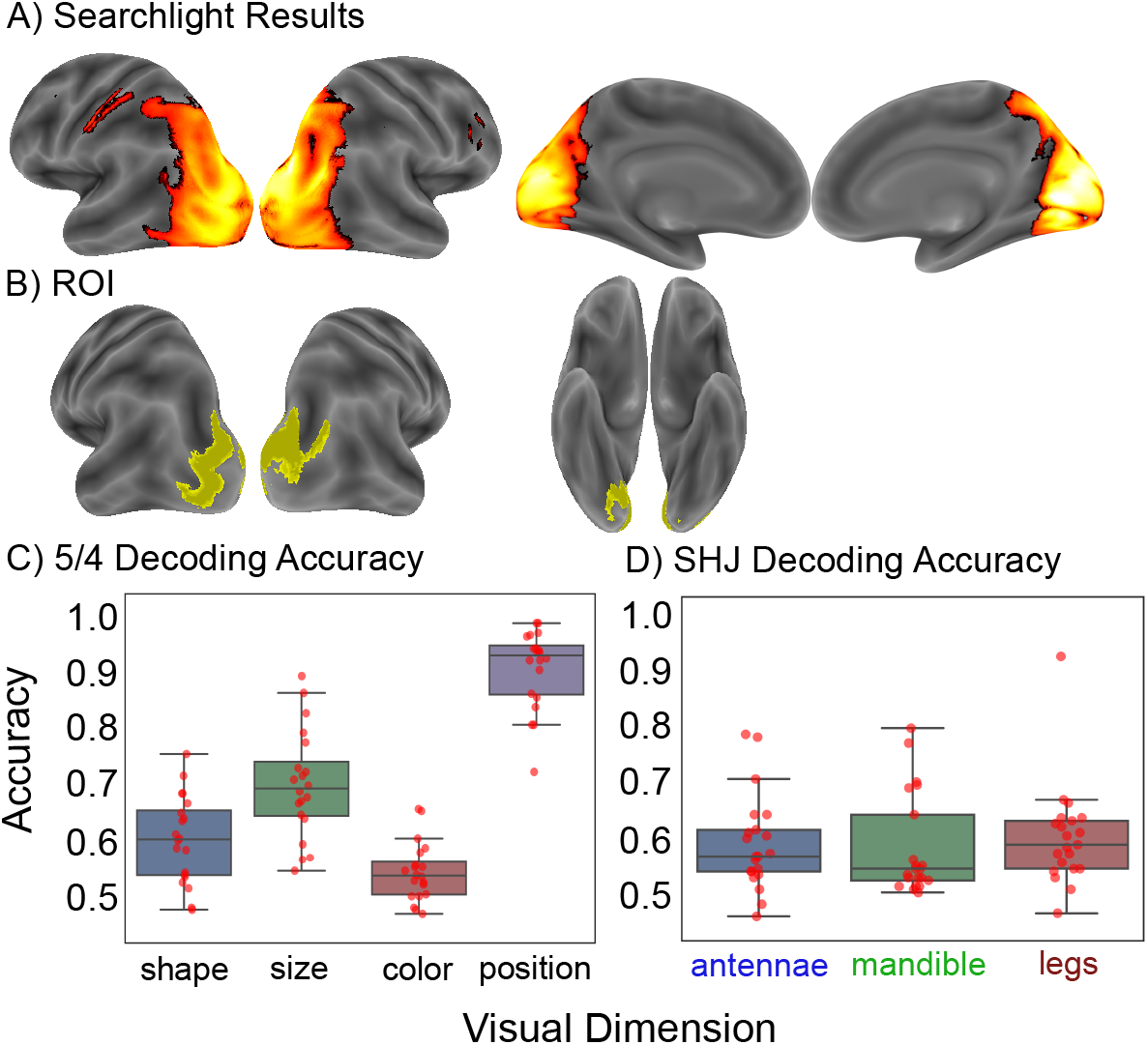
**A**) For the “5/4” dataset, a searchlight analysis indicated that binary perceptual dimensions could be decoded from widespread visual regions (including occipital, temporal and parietal cortex), right inferior frontal sulcus, and left post-central motor cortex (the family-wise error rate was controlled at the voxel-level *p* < 0.001). **B**) To isolate voxels most strongly representing the stimulus features, we raised the statistical threshold, resulting in the ROI illustrated in yellow. **C**) “5/4” Dataset Binary Feature Decoding. Red dots indicate scores from individual participants. **D**) SHJ Binary Feature Decoding. The same ROI (B) was used in both datasets.

SUSTAIN was initialized with no clusters, and with equivalent weights assigned to each stimulus dimension. Its learning parameters were first fit to the learning performance of each participant using a maximum-likelihood genetic algorithm procedure. The model was fit in such a way that, after learning one problem, the model state was used as the initial state for the subsequent problem. In this way, the model was fit under the assumption that learning of one task would influence later behavior. Once the learning parameters of the model were optimized, they were fixed, and the attentional parameters were extracted from the second two runs of each task (in which learning had largely stabilized). This yielded distinct sets of attentional parameters for each participant and each task. More information about the model can be found in appendix C.

### Image Processing

Preprocessing included motion correction, and coregistration of the anatomical images to the mean of the functional images (using Statistical Parametric Mapping (SPM), version 6470). All MVPA analyses were performed in native space without smoothing. For group-level analyses, the statistical maps from each participant were warped to Montreal Neurological Institute (MNI) atlas space using Advanced Normalization Tools (ANTs; Avants et al., 2009), and then smoothed with a 6mm full-width at half maximum Gaussian kernel. The ROI derived from group-level analyses were transformed back into each participants native space for ROI-level analyses. We performed MVPA on the unsmoothed, single-trial, *t*-statistic images (Misaki et al., 2010) derived from the least-squares separate procedure (LSS; Mumford et al., 2012). We used SPM to estimate the LSS images for the “5/4” dataset, but used the NiPy python package (http://nipy.org/nipy/index.html) for the SHJ dataset, as it tends to run more efficiently, and this study used a multiband sequence with smaller voxel dimensions.

## Results

### Representations of Individual Visual Features

To identify regions most strongly representing the stimulus features, we performed a cross-validated searchlight analysis (sphere radius = 10mm.; Kriegeskorte et al., 2006)^2^ in which we decoded each of the four visual features (position, shape, color and size). We performed the analysis in native anatomical space, using a linear support vector classifier (SVC, C=0.1; using the Scikit-learn python package; Pedregosa et al., 2011) in conjunction with a five-fold, leave-one-run out, cross-validated procedure. This involved repeatedly training the model on four of the five runs, and testing whether it could accurately predict the stimulus features associated with the held-out neuroimaging data.

After centering each of the resultant statistical maps at chance (50% for each visual feature), we created a single map for each participant, which reflected the average, above chance, decoding accuracy across features. We then normalized each map to to MNI space and, in order to identify regions supporting above chance feature decoding, performed a group-level permutation test. This involved randomly flipping the sign of the statistical maps 10,000 times (using the randomise function from the Oxford Centre for Functional MRI of the Brain Software Library (FSL); Winkler et al., 2014). The familywise error rate was controlled using a voxelwise threshold of *p* < 0.001. This identified right middle frontal gyrus (BA9) and left post-central motor cortex, as well as widespread visual and association cortex, extending dorsally from occipital pole to the bilateral superior extrastriate cortex and bilateral intraparietal sulcus (IPS), and ventrally into the bilateral lingual gyrus (Table 1). As this procedure yielded a diffuse pattern of spatial activity, we increased the minimum *t*-statistic threshold (from 6.24 to 9) to isolate voxels most strongly representing the individual stimulus features. This removed voxels belonging to the bilateral inferior occipital cortex, left lingual gyrus, bilateral intraparietal sulcus, and bilateral precuneus. The resultant ROI is illustrated in Figure 3.B.

### Effects Associated with Conceptual Knowledge “5/4” Dataset

First, we confirmed that each stimulus feature could be decoded significantly above chance from the ROI illustrated in Figure 3.B. Although estimating effect sizes on voxels selected through non-orthogonal criteria is circular, testing significance at the ROI-level has been recommended to confirm that information exists, not only at the level of the searchlight sphere, but also at the level of the ROI (Etzel et al., 2013). This analysis also allows us to illustrate the individual feature decoding accuracies for each participant (Figure 3.C). The analyses were performed in the native anatomical space of each participant using the cross-validated SVC analysis described above (but setting the C parameter to 1 instead of 0.1, which was chosen for the searchlight analysis to improve computational efficiency).^3^ Each feature could be decoded at rates significantly above chance (shape: *M* = 0.60, *SE* = 0.02, t(19) = 5.78, *p* < 0.001, size: *M* = 0.70, *SE* = 0.02, t(19) = 9.29, *p* < 0.001, color: *M* = 0.54, *SE* = 0.01, t(19) = 3.64, *p* = 0.002, position: *M* = 0.91, *SE* = 0.02, t(19) = 25.96, *p* < 0.001).

Next, we investigated whether the decoding accuracy of the individual perceptual dimensions covaried with the GCM attentional parameters. To do so, we fit a mixed-effects linear regression analysis (as implemented in the lme4 package for R) using restricted maximum likelihood (ReML). We included fixed-effects terms for the intercept, the attentional weights, and each visual dimension (e.g., “color”). We also included random effects terms (which were free to vary between participants) for the intercept and the attention weight parameters. This allowed us to control for baseline differences in decoding accuracy between participants, and for shared (group-level) differences in decoding accuracy between visual dimensions. We used the Kenward-Roger approximation (Kenward & Roger, 1997) to estimate degrees of freedom (reported below), and used single-sample *t*-tests to calculate p-values for each coefficient (using the pbkrtest package for R; Halekoh & Højsgaard, 2014)^4^. We computed 95% confidence intervals using bootstrap resampling (1000 simulations). The decoding accuracy of each stimulus dimension positively covaried with the behaviorally-derived GCM parameters (*b* = 0.08, 95% CI = [0.01, 0.16], *SE* = 0.04, t(28.71) = 2.26, *p* = 0.032), indicating that the decoding accuracy of these representations reflected their importance during decision-making.

To investigate the sensitivity of occipitotemporal feature representations to individual differences in GCM attentional weights, we conducted a permutation test. This involved shuffling the attentional weight parameters between participants (i.e., swapping the weights derived from one participant with those derived from another), and repeating the regression analysis (described above) 10,000 times. On each permutation, the correspondence for category dimensions (i.e., the dimensions depicted in Table 2, as opposed to the stimulus dimensions illustrated in Figure 1) was preserved, such that the dimensional weights derived from the behavior of one participant were assigned to the same dimensions, but to a different participant.

The unpermuted beta coefficient (b = 0.08) was significantly greater than those composing the null distribution (P = 0.994), indicating that the decoding accuracy of the occipitotemporal representations was sensitive to between-subject differences in the attentional weights. This could reflect idiosyncratic differences in behavioral strategy, and/or effects associated with perceptual saliency. Therefore, to investigate whether visual salience may have influenced attention, we conducted a repeated measures ANOVA for the perceptual features. There was no significant relationship between these visual features and the attentional parameters (F(3,57) = 0.68, *p* = 0.56). A Bayesian repeated-measures ANOVA (Rouder et al., 2017), additionally indicated that the null model was 4.65 times more likely than the alternative hypothesis. These results provide evidence that the observed effects were not driven by visual characteristics of the stimulus features.

### SHJ Dataset

First, we confirmed that each stimulus feature could be decoded significantly above chance from the ROI illustrated in Figure 3.B. Using a four-fold, leave-one-run out cross-validation strategy, we used a linear support vector classifier (C=1) to decode each visual feature across all runs (including both early and late learning epochs), retaining only estimates for the last two runs (which corresponded to the late-learning phase in which behavior had largely stabilized). This four-fold cross-validation strategy yielded better decoding accuracy than a two-fold approach based on only the last two runs. This improvement reflects the increased amount of training data available in the 4-fold approach, and suggests that the multivariate patterns reflecting the individual visual features were stable across learning. Each feature could be decoded at rates significantly above chance (Figure 3.D; antennae: *M* = 0.57, t(20) = 3.82, *p* = 0.001; mandibles: *M* = 0.56, *t*(20) = 3.22, *p* = 0.004; legs: *M* = 0.58, t(20) = 4.17, *p* < 0.001).

Next, we investigated whether the decoding accuracy associated with the features covaried with SUSTAIN’s attentional parameters. To do so, we used a mixed-effects linear regression analysis to predict decoding accuracy from attention weight, visual dimension, run and rule. As described in the Methods section, distinct attentional weights were derived for each subject and each rule. The decoding accuracy for each separate run was included in the analysis. The model included fixed-effects parameters for these four variables, and random-effects parameters for the intercept, attention weight, and run (which were free to vary by participant). This allowed us to control for differences in decoding accuracy across visual dimensions and participants (as with the model used for the “5/4” dataset), while additionally controlling for effects of rule and idiosyncratic differences in behavioral performance during the last two runs. Mirroring the findings from the “5/4” dataset, we found that the decoding accuracy of these patterns positively covaried with the attention parameters derived from SUSTAIN (b = 0.09, 95% CI = [0.004, 0.17], *SE* = 0.04, *t*(61) = 2.13, *p* = 0.038).

To investigate the sensitivity of occipitotemporal feature representations to individual differences in SUSTAIN’s attentional parameters, we conducted a permutation test similar to that described above (i.e., for the “5/4” experiment). This involved shuffling the attentional weight parameters between participants 10,000 times (preserving the correspondence for both rule and abstract feature). This means that the attentional weight derived from the behavior of one participant, for one particular rule and one particular category feature, was assigned to the same rule and feature, but to a different participant. The slope parameter associated with the unpermuted data (b = 0.09) was significantly greater than those composing the permuted null distribution (P = 0.979), suggesting that the visual feature representations were sensitive to idiosyncratic differences in attentional weights. A repeated measures ANOVA indicated that the perceptual dimensions did not influence the attentional parameters (F(2,44) = 1.27, *p* = 0.291). A Bayesian repeated measures ANOVA additionally indicated that the null model was 1.98 times more likely than the alternative hypothesis, providing evidence that the attentional weights were not influenced by visual properties of the stimulus features.

## Discussion

Although differing substantially in how concepts are represented (e.g., as exemplars, prototypes, or clusters), formal categorization theories (e.g., Love et al., 2004; Kruschke, 1992; Nosofsky, 1986) tend to share a similar conception of selective attention. In these models, conceptual knowledge contorts multidimensional psychological space such that differences along behaviorally-relevant dimensions are accentuated, and differences along irrelevant dimensions are down-weighted (Figure 1, and Equations 1 & 4 in appendix C). In two datasets (Mack et al., 2013, 2016), we evaluated the neurobiological plausibility of this idea by investigating whether occipitotemporal stimulus feature representations covaried with attention parameters derived from formal categorization models. We found that this effect was not only apparent at the group-level, but was sufficiently sensitive to reflect individual differences in conceptual knowledge.

Several previous studies have demonstrated that occipitotemporal stimulus representations are modulated by selective attention (e.g., Buffalo et al., 2010; Jehee et al., 2011; Kamitani & Tong, 2005, 2006; Luck et al., 1997; Motter, 1993; Reynolds et al., 2000; Reynolds & Chelazzi, 2004) and by learned conceptual knowledge (e.g., Folstein et al., 2013; Li et al., 2007; Sigala & Logothetis, 2002). These studies have relied on statistically-powerful contrastive approaches, in which representations of attended stimulus dimensions are compared to those of unattended dimensions. A known effect is that neural signatures of attended stimulus dimensions are more discriminable than those that are unattended. This implies that occipitotemporal representational space might resemble that conceptualized by formal categorization theory (e.g., Nosofsky, 1986; Love et al., 2004; Kruschke, 1992). Specifically, the expansion and contraction of this space might reflect individual differences in the importance assigned to each stimulus dimension. However, as the contrastive approach defines selective attention with regards to the experimental paradigm, it is insensitive to individual differences in categorization strategy (e.g., Craig & Lewandowsky, 2012; Little & McDaniel, 2015; McDaniel et al., 2014; Raijmakers et al., 2014). Here, we link individual differences in behavior to individual differences in neural representation through consideration of the attentional parameters derived from formal categorization models.

We are not the first to link brain and behavior via latent model parameters. In the perceptual decision-making literature, for instance, several groups have fit the drift diffusion model (Ratcliff, 1978) to behavioral data, and identified regions of the brain where the BOLD response reflects variation in its drift rate, bias, and threshold parameters (e.g., Mulder et al., 2012; Forstmann et al., 2008; Purcell et al., 2010). As in the present study, several of these studies demonstrated that individual differences in behavioral strategy are reflected in the brain. Instead of linking latent model parameters to univariate BOLD amplitude, however, we used MVPA to link latent parameters to multivoxel representations of the stimulus features. This provided a precise test of the idea that selective attention contorts neural representational space.

These endogenous attentional effects are thought to arise through communication with other areas of the brain. In lateral frontal cortex, for instance, effects of endogenous attention occur earlier in time than in occipitotemporal cortex (Bichot et al., 2015; Baldauf & Desimone, 2014; Zhou & Desimone, 2011). Inactivation of these frontal regions (e.g., ventral prearcuate sulcus, or entire lateral prefrontal cortex) has also been associated wtih a reduction in the magnitude of attentional effects in occipitotemporal cortex (Bichot et al., 2015; Gregoriou et al., 2014). Interestingly, contextually-sensitive effects of endogenous attention have also been observed in the lateral geniculate nucleus (LGN), suggesting that some aspects of attention precede those in cortex (O’Connor et al., 2002; McAlonan et al., 2008; Saalmann & Kastner, 2011).

Finally, although we observed effects of selective attention across two different stimulus sets (abstract shapes in the “5/4” experiment, and insects in the SHJ experiment), and across multiple category structures (the “5/4” problem described by Medin & Schaffer (1978), and the Type I, II and VI problems described by Shepard et al. (1961)), these effects might not be apparent for all stimuli and tasks. For instance, although category training can improve perceptual discriminability of relevant stimulus features when stimuli consist of perceptually-separable features (Garner, 1976), this may not occur for integral dimensions (Op de Beeck et al., 2003) or for stimuli defined according to “blended” stimulus morphspaces (Folstein et al., 2013). More work is needed to better understand how attention influences occipitotemporal representations for such stimuli. One possibility is that selective attention does not warp *perceptual* representations of integral stimulus dimensions, but might operate on abstract cognitive or “decisional” representations in higher-order cortex (Jiang et al., 2007; Nosofsky, 1987).

### Conclusions

Category training is known to induce changes in both perceptual (Folstein et al., 2012; Op de Beeck et al., 2003; Goldstone et al., 1996; Goldstone, 1994; Gureckis & Goldstone, 2008) and neural sensitivity (e.g., Dieciuc et al., 2017; Folstein et al., 2013, 2015; Li et al., 2007; Sigala & Logothetis, 2002). In two datasets, we demonstrate that occipitotemporal stimulus representations covary with the attentional parameters derived from formal categorization theory. This effect was sufficiently sensitive to reflect individual differences in conceptual knowledge, which implies that these occipitotemporal representations are embedded within a space closely resembling that predicted by formal categorization theory (e.g., Nosofsky, 1986; Love et al., 2004; Kruschke, 1992).

By linking brain and behavior through the latent attentional parameters of cognitive models, we also link two (somewhat) disparate literatures. In the neuroscience literature, effects of selective attention are typically examined using highly-structured decision problems, and selective attention is investigated by contrasting different aspects of the experimental design (i.e., relevant vs. irrelevant stimulus dimensions). In the cognitive categorization literature, researchers have focused on developing models that accurately account for behavioral patterns of generalization across different goals and tasks. Our results indicate that these cognitive models can be used to examine effects of selective attention in the brain. This is the case, even for ill-defined decision problems (such as the “5/4” task), as the models are able to successfully account for individual differences in conceptual knowledge.

## Appendix A: Searchlight Results

**Table 1:** “5/4” Dataset Binary Feature Decoding: Searchlight Results. The familywise error rate was controlled at the voxel level (*p* < 0.001)

| Size | x | y | z | $t$ | BA | Region |
| --- | --- | --- | --- | --- | --- | --- |
| 23972 | 14 | -74 | 4 | 12.717 | 18 | Calcarine_R |
|  | 28 | -62 | 54 | 10.5 | 7 | Parietal_Sup_R |
|  | -54 | -18 | 44 | 7.256 | 3 | Postcentral_L |
| 233 | 50 | 24 | 26 | 7.447 | 48 | Frontal_Inf_Tri_R |
| 22 | -46 | 12 | 44 | 6.621 | 9 | Frontal_Mid_L |

## Appendix B: Category Structures

**Table 2:** The “5/4” Category Structure. Prior to scanning, participants learned, through trial and error, to categorize the first nine stimuli (category “A”: A1-A5; category “B”: B1-B4) illustrated in Figure 2.A. During scanning, they categorized both the training and the transfer (T1-T7) stimuli. Perceptual stimulus dimensions (Figure 2) were pseudo-randomly assigned to category dimensions for each participant.

| Stimulus | D1 | D2 | D3 | D4 |
| --- | --- | --- | --- | --- |
| A1 | 1 | 0 | 0 | 0 |
| A2 | 1 | 0 | 1 | 0 |
| A3 | 0 | 1 | 0 | 0 |
| A4 | 0 | 0 | 1 | 0 |
| A5 | 0 | 0 | 0 | 1 |
| B1 | 1 | 1 | 0 | 0 |
| B2 | 1 | 0 | 0 | 1 |
| B3 | 0 | 1 | 1 | 1 |
| B4 | 1 | 1 | 1 | 1 |
| T1 | 0 | 1 | 1 | 0 |
| T2 | 1 | 1 | 1 | 0 |
| T3 | 0 | 0 | 0 | 0 |
| T4 | 1 | 1 | 0 | 1 |
| T5 | 0 | 1 | 0 | 1 |
| T6 | 0 | 0 | 1 | 1 |
| T7 | 1 | 0 | 1 | 1 |

**Table 3:** SHJ Category Structures. Participants learned by trial-and-error to perform the type I (a one-dimensional rule-based categorization task), type II (a two-dimensional XOR rule-based categorization task), and type VI (a three-dimensional task requiring memorization of the individual stimuli) problems during scanning. For each participant, perceptual stimulus dimensions (Figure 2) were randomly assigned to these abstract category dimensions.

| Stimulus | D1 | D2 | D3 | Type1 | Type2 | Type6 |
| --- | --- | --- | --- | --- | --- | --- |
| 1 | 0 | 0 | 0 | A | A | A |
| 2 | 0 | 0 | 1 | A | B | B |
| 3 | 0 | 1 | 0 | A | B | B |
| 4 | 0 | 1 | 1 | A | A | A |
| 5 | 1 | 0 | 0 | B | A | B |
| 6 | 1 | 0 | 1 | B | B | A |
| 7 | 1 | 1 | 0 | B | B | A |
| 8 | 1 | 1 | 1 | B | A | B |

## Appendix C: Computational Models and Attentional Parameters

For the first dataset (Mack et al., 2013), we considered the Generalized Context Model (GCM; Nosofsky, 1986), which posits that conceptual knowledge consists of memory for individual exemplars. For the SHJ experiment (Mack et al., 2016), we considered the attentional parameters from SUSTAIN (Supervised and Unsupervised STratified Adaptive Incremental Network; Love et al., 2004). Details about the models can be found in the original papers. Here, we provide a brief overview of each.

### GCM

In the Generalized Context Model (GCM; Nosofsky, 1987), the psychological distance, *d* between stimuli *i* and *j* can be calculated as the attentionally-weighted sum of their unsigned differences across dimensions, *k*:

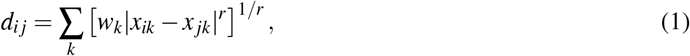

where *w* indicate the attentional parameters assigned to each dimension. The *r* parameter is set to 1 (city-block distance) for perceptually separable stimulus dimensions (as in the “5/4” dataset), and *r* is set to 2 (Euclidean distance) for integral dimensions (Garner, 1976). Similarity is an exponentially-decaying function of psychological distance:

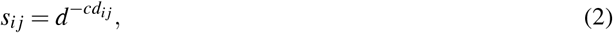

where the shape of the similarity gradient is influenced by the sensitivity parameter, *c*. The probability of choosing category “A”, given stimulus, *i*, is given by the choice rule:

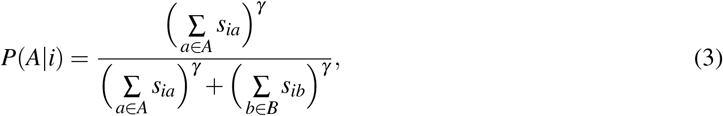

where *γ* governs the degree of deterministic responding.

### SUSTAIN

SUSTAIN is a semi-supervised clustering model, which incrementally learns to solve categorization problems by first applying simple solutions, and then increasing complexity as required. Through experience, the model can learn to group similar items into common clusters, and can make inferences about novel stimuli based on its perceptual similarity to existing clusters (i.e., based on perceptual similarity, clusters compete to predict latent stimulus attributes). When unexpected feedback is received, the model can also learn in a supervised fashion by creating a new cluster to represent the novel stimulus.

In SUSTAIN, all clusters contain receptive fields (RF’s) for each stimulus dimension. As new stimuli are added to the cluster, the model learns by adjusting the position of each RF to best match the cluster’s expectation for novel stimuli. As the RF is an exponential function, a cluster’s activation, *α*, decreases exponentially with distance from its preferred value:

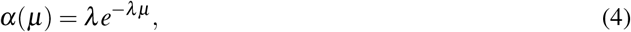

where *μ* represents the distance of the stimulus dimension value from the cluster’s preferred stimulus dimension value, and where *λ* represents the tuning (or width) of the RF. The *λ* parameters are specific to dimensions, but are shared across dimensions, and so, like the attentional parameters in the GCM, the *λ* parameters in SUSTAIN modulate the influence of each stimulus dimension on the overall decision outcome.

The overall activation of a cluster, *H*, involves consideration of each dimension, *k*:

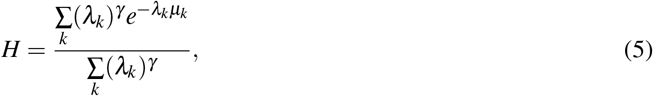

where the *γ* parameter (which is always non-negative) modulates the influence of the *λ* parameters on the choice outcome. When *γ* is large, attended dimensions (which are associated with large *λ* values, and narrow RF’s), dominate the activation function (eq. 5); when 7 is zero, the *λ* parameters are ignored, and all dimensions exert equal influence on the choice.

SUSTAIN was fit to the SHJ dataset in a supervised fashion, using the same trial order experienced by the participants; it was also fit across rule-switches, such that learning from one task was carried over to the next. Thus, SUSTAIN was capable of reflecting learning, as well as carry-over effects associated with previously learned rules.

## Footnotes

1 In their paper, Mack et al. (2016) focus on effects associated with the type I and type II rules.

2 This involves moving an imaginary sphere throughout the brain; repeatedly investigating how well the voxels within the sphere can decode a variable of interest.

3 The C parameter modulates the penalty associated with training error. With large values, the classifier will choose a small-margin hyperplane, and training accuracy will be high. With smaller values, out-of-sample performance is often improved, but more training samples may be misclassified. C=1 is a common default setting for fMRI.

4 This provides a more conservative test than the likelihood ratio test or the Wald approximation (Luke, 2016).

## Acknowledgments

We thank the authors of the original studies for sharing their data. This work was supported by NIH Grant 1P01HD080679, Leverhulme Trust grant RPG-2014-075 and Wellcome Trust Senior Investigator Award WT106931MA to BCL.

